# Size and position dependent cytoplasm viscoelasticity through hydrodynamic interactions with the cell surface

**DOI:** 10.1101/2022.09.27.509722

**Authors:** Javad Najafi, Serge Dmitrieff, Nicolas Minc

## Abstract

Many studies of cytoplasm rheology have focused on small components in the sub-micrometer scale. However, the cytoplasm also baths large organelles like nuclei, microtubule asters or spindles that often take significant portions of cells and move across the cytoplasm to regulate cell division or polarization. Here, we translated passive components of sizes ranging from few up to ~50 percent of the cell diameter, through the vast cytoplasm of live sea urchin eggs, with calibrated magnetic forces. Creep and relaxation responses indicate that for objects larger than the micron size, the cytoplasm behaves as a Jeffreys’ material, viscoelastic at short time-scales and fluidizing at longer times. However, as components size approached that of cells, cytoplasm viscoelastic resistance increased in a non-monotonic manner. Flow analysis and simulations suggest that this size-dependent viscoelasticity emerges from hydrodynamic interactions between the moving object and the static cell surface. This effect also yields to position-dependent viscoelasticity with objects initially closer to the cell surface being harder to displace. These findings suggest that the cytoplasm hydrodynamically couples large organelles to the cell surface to restrain their motion, with important implications for cell shape sensing and cellular organization.

**Significance Statement:** Large-sized organelles like nuclei or mitotic spindles typically translocate through the cytoplasm to regulate cell division or polarity, but their frictional interaction with the cytoplasm and the cell surface remain poorly addressed. We used *in vivo* magnetic tweezers, to move passive components in a range of size in the cytoplasm of living cells. We found that the mobility of objects with sizes approaching that of cells, can be largely reduced as a result of hydrodynamic interactions that couple objects and the cell surface through the cytoplasm fluid.

## INTRODUCTION

The cytoplasm is a complex heterogenous medium crowded with macromolecules and entangled cytoskeleton networks (1). These define rheological properties that may influence molecular processes ranging from molecular diffusion to reaction kinetics and protein folding (2–5). However, the cytoplasm also hosts the motion of much larger elements closer to the cellular scale, for which the impact of cytoplasm properties remain much less understood. For instance, during critical events such as fertilization, cell polarization or asymmetric divisions, cells actively displace large nuclei, microtubule asters or mitotic spindles across their cytoplasm (6, 7). These organelles are moved by forces generated from active polar cytoskeletal networks such as actomyosin bundles or microtubules and associated motors, that need to overcome mechanical resistance from the cytoplasm medium (8). Therefore, addressing the nature and magnitude of frictional interactions of large components with the cytoplasm remains an important endeavor for cellular spatial organization.

Many past studies have measured rheological properties of bulk cytoplasm, using either extracts *in vitro*, or by actuating endogenous or foreign probes in living cells. These have shown that cytoplasm response will depend on time-scale, force amplitude or component size (9–13). The question of object size has notably raised important notions for cytoplasm mechanics, as for components smaller or larger than a typical mesh size, the cytoplasm may exhibit different rheological signatures ranging from fluid, to viscoelastic, poroelastic, or even glassy materials (14–19). However, these studies were restricted to regimes of relatively small objects typically below the micron size, as well as low forces and displacement amplitudes. As objects reach closer to the cell size and move across longer distances, they are predicted to drag and shear large portions of the cytoplasm fluid (20–22). In such regime, cytoplasm resistance may be influenced by boundary conditions at the static cell surface, through so-called wall effects. These have been well documented in fluid mechanics, and emerge from hydrodynamic interactions that couple a moving object with a static wall. They are predicted to enhance frictional coefficients by a significant amount if the object contained by the walls approaches the size of the container (23). To date, however, direct evidence that these effects are relevant to living cells is still lacking, in part given the technical difficulty of applying calibrated forces onto large components *in vivo*.

Here, we employed calibrated *in vivo* magnetic tweezers to move passive components of sizes ranging from ~1% to ~50% of the cell diameter across the cytoplasm of living cells. By tracking resultant cytoplasm shear flows and using finite element hydrodynamic simulations, we demonstrate how object size disproportionally restrain its mobility as a result of hydrodynamic interactions with the cell surface. We also find that these interactions can yield to position-dependent cytoplasm resistance with objects initially closer to the cell surface becoming harder to pull or push. This work suggests that large organelles may be coupled to the cell surface without any direct cytoskeletal connections, and highlights the underappreciated impact of confinement by cell boundaries to organelle motion.

## RESULTS

### Probing bulk cytoplasm rheology at multiple length scales

We sought to probe the rheology of bulk cytoplasm in response to the typical translational motion of relatively large organelles, such as *e.g*. micrometric acidic organelles or even larger nuclei or mitotic spindles. These objects are commonly moved by molecular motors with directional speeds ranging from fractions up to tens of μm/s, across distances in the scale of few to few tens of micron (24). We used sea urchin unfertilized eggs as model cell types. These are large cells, with a diameter of 95 μm, amenable to quantitative injection, that are arrested in a G0- or G1-like state of the cell cycle before fertilization (25). Importantly, in contrast to fertilized eggs, unfertilized eggs do not feature any large-scale cytoskeletal organization, with F-actin and microtubule filaments distributed throughout the cytoplasm, and no detectable thick actin cortex (Fig. S1A-B) (26). In addition, Particle Imaging Velocimetry (PIV) of relatively large granules, did not reveal any persistent cytoplasm flows, suggesting that the cytoplasm material may be considered at rest in these cells (Fig. S1C).

To compute the viscoelastic response of the cytoplasm at different length-scales, we injected single magnetic beads of 1 μm in diameter, as well as bead aggregates of variant sizes from few up to ten microns (27, 28). To reach component sizes closer to cell size, we injected a suspension of soya oil mixed with hydrophobic ~1 μm magnetic beads inside eggs. Upon injection, this suspension formed large magnetized oil droplets that ranged in size from 18 to 45 μm (corresponding to ~20-48% of the cell diameter) (20). We applied calibrated forces, by approaching a magnet tip close to the cell surface. This caused components to translate through the cytoplasm along the magnetic gradient (Fig. 1A) (28). Magnetic forces and their dependence on bead type, magnet-bead distance and aggregate size were calibrated *in vitro* using test viscous fluids (Figs. S1D-F).

**Figure 1.**
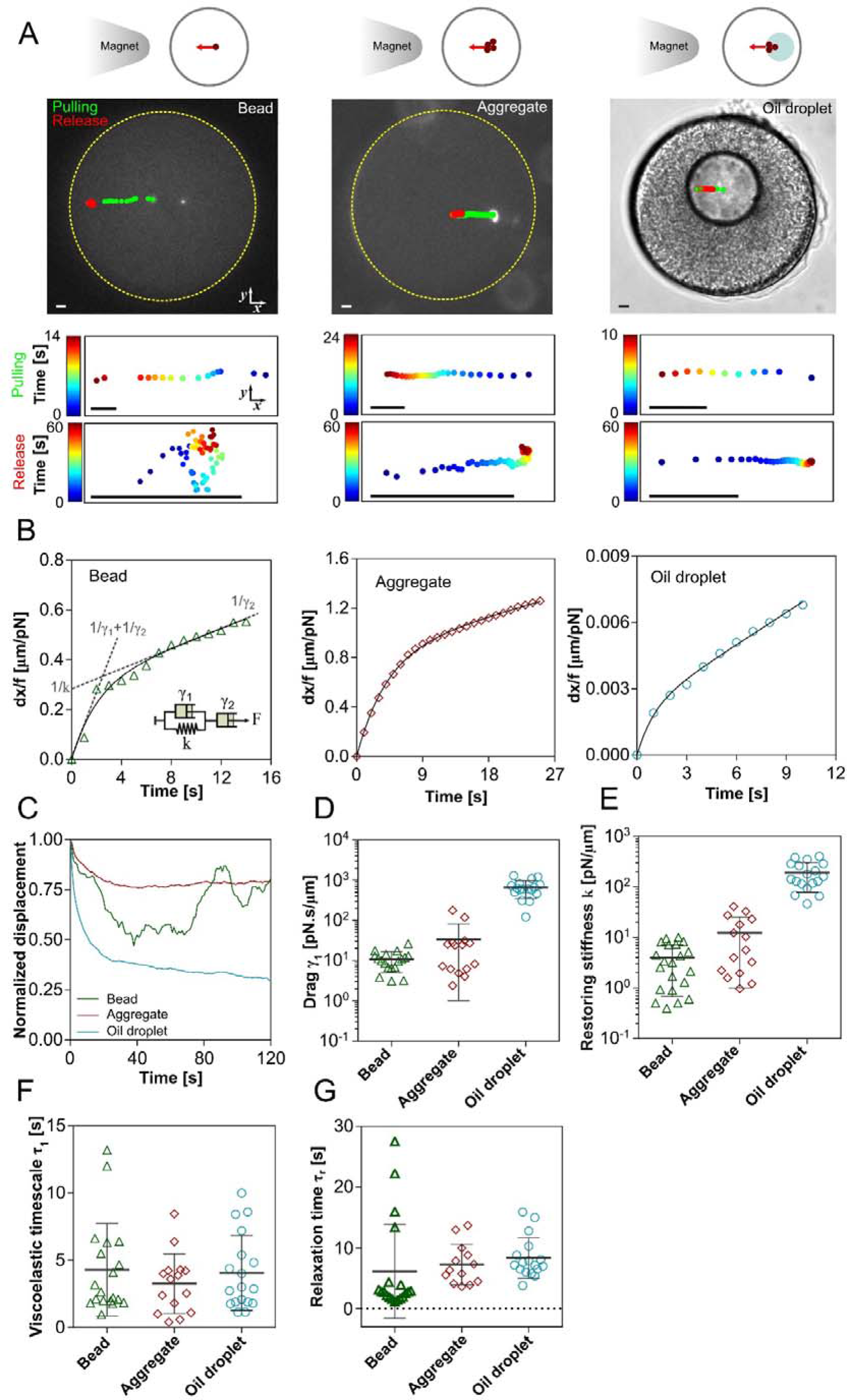
Probing cytoplasm rheology at large length-scales. (A) Sea urchin unfertilized eggs injected with single 1 μm beads, beads aggregate with sizes ranging from ~4-12 μm and large oil droplets containing hydrophobic beads. Calibrated magnetic forces were applied to translate these magnetized objects in the cytoplasm. The trajectories of the objects tracked at 1 Hz during force application (green) and force release (red) are overlaid on the cells, and also represented in time color coded graphs at the bottom of cells. (B) Representative displacement curves scaled by forces plotted as a function of time for beads, aggregates and magnetized oil droplets. The Jeffreys’ viscoelastic model is depicted as an inset in the graph for single beads, with representative constants overlaid on the curve. Solid lines are fits of the Jeffreys’ model. (C) Normalized recoiling displacements plotted as a function of time for the same objects as in B. Note that the recoil of small beads towards their initial position is non-directional due to intrinsic cytoplasm noise. (D-G) Cytoplasm viscoelastic parameters for objects of variant sizes computed by fitting creep and relaxation curves with Jeffreys’ model (n= 21, 15 and 18, respectively): viscous drags (D), restoring stiffness (E), viscoelastic time-scale during the rising phase (F) and in the releasing phase (G). Error bars correspond to +/− SD. Scale bars, 5μm.

Interestingly, all these objects exhibited similar creep responses to an applied force. First, objects moved at constant speed, indicating a viscous behavior. Then the displacement-time curve inflected reflecting elastic responses. At longer time-scales the behavior was linear again indicating a fluidization of elastic elements (20, 27, 29) (Fig. 1B). Accordingly, when the force was released, objects recoiled back towards their initial positions with partial recoils that reflected a dissipation of stored elastic energy. However, during recoils, small beads were more prone to random forces and space exploration, exhibiting a much less directional recoiling behavior (Fig. 1C and Movie S1).

To quantify cytoplasm viscoelastic parameters, we fitted both creep and recovery curves with a 1D Jeffreys’ model, consisting of a Kelvin-Voigt viscoelastic element in series with a dashpot which represents the dissipation of the material in the long asymptotic time (27, 30) (Fig. 1B inset). This allowed to compute a short-term viscous drag of the translating object, γ_1_, a restoring stiffness k, and thus the viscoelastic relaxation time-scale of cytoplasm material τ_1_ = γ_1_/k. Mean values of viscous drags increased with object size, from 11 ± 6 pN.s/μm (mean ± std) for small 1 μm beads, to 33 ± 49 pN.s/μm for larger aggregates and up to 660 ± 312 pN.s/μm for very large oil droplets (Fig. 1D). Similarly, the restoring stiffness increased from 4 ± 3 pN/μm, to 12 ± 13 pN/μm and 191 ± 112 pN/μm, respectively (Fig. 1E). Importantly, however, the relaxation time-scales was mostly independent of object size, and was around ~4 s during force loading and ~7 s during relaxation (Figs. 1F-G). These results suggest similar general rheological responses of the cytoplasm to translating objects ranging in size from few up to 50% of cell size.

To confirm that viscoelastic responses and parameters reflected bulk cytoplasm properties, we next injected magnetic droplets and rinsed eggs in diluted (50%) or concentrated (120%) artificial sea water (ASW) to vary extracellular osmolarity and, respectfully decrease or increase macromolecular crowding (3, 31, 32) (Figs. 2A-B and Movie S2). Both creep and relaxation responses had similar shapes as in controls, but viscous drag and restoring stiffness values were markedly different (Fig. 2C). In hypoosmotic conditions (50% ASW), they were reduced by ~4.2X and 3.8X, respectively, and increased by 1.6 and 2.8X in hyperosmotic conditions (120% ASW). Accordingly, characteristic viscoelastic time-scales during the pulling phase were similar in control and hypoosmotic conditions (4.1 ± 2.8 s and 3.9 ± 2.4 s), but slightly reduced to 2.2 ± 1.5 s in hyperosmotic conditions. These results support that the measured viscoelastic resistance to these large translating objects directly relate to cytoplasm crowding and mechanical properties.

**Figure 2.**
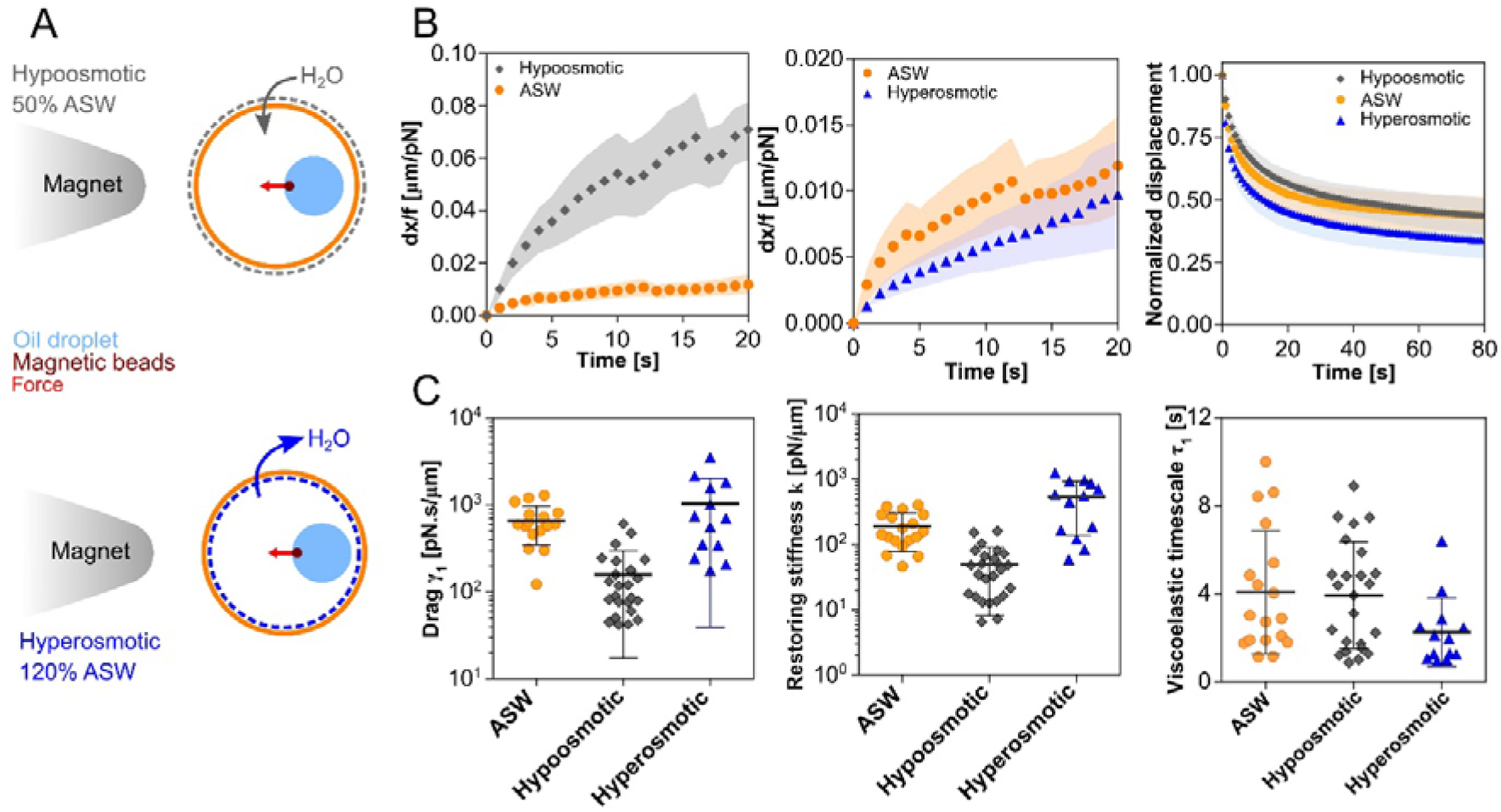
Viscoelastic properties depend on cytoplasm crowding. (A) Schematic of experiments in which cytoplasm crowding is modified by placing cells in hypoosmotic or hyperosmotic Artificial Sea Water (ASW). (B) Displacement curves scaled by forces during pulling phases, and normalized relaxation curves after force retraction for control (n=18), hypoosmotic (n=26), and hyperosmotic (n=13) conditions for oil droplets. (C) Viscous drag, restoring stiffness, and viscoelastic time-scale in the rising phase at different osmotic conditions. Shaded areas represent +/− SEM and error bars correspond to +/− SD.

### Cytoplasm viscoelastic shear flows associated to the motion of large objects

To understand how the motion of different sized objects impact cytoplasm organization, we mapped cytoplasm flows, by tracking granules visible in bright field with PIV. The movement of small beads, did not create notable large-scale flows, reflecting the high viscosity of the cytoplasm that screens perturbations at long length-scales (Fig. S2A). In contrast, the motion of very large droplets, yielded notable and reproducible cytoplasm flow fields. The maximum cytoplasm flow speed was on the order of droplet speed at the front and back of the droplet along the pulling direction and dropped rapidly away from the droplet. Moreover, two vortices were generated at the upper and lower parts of the pulling axis, often asymmetric in size when the droplet and force axis were, for instance, off–center (Figs. 3B-D and Movie S3). This suggests that large translating objects create significant shear in the cytoplasm with magnitudes and organization that may depend on object size and its distance to cell boundaries.

**Figure 3.**
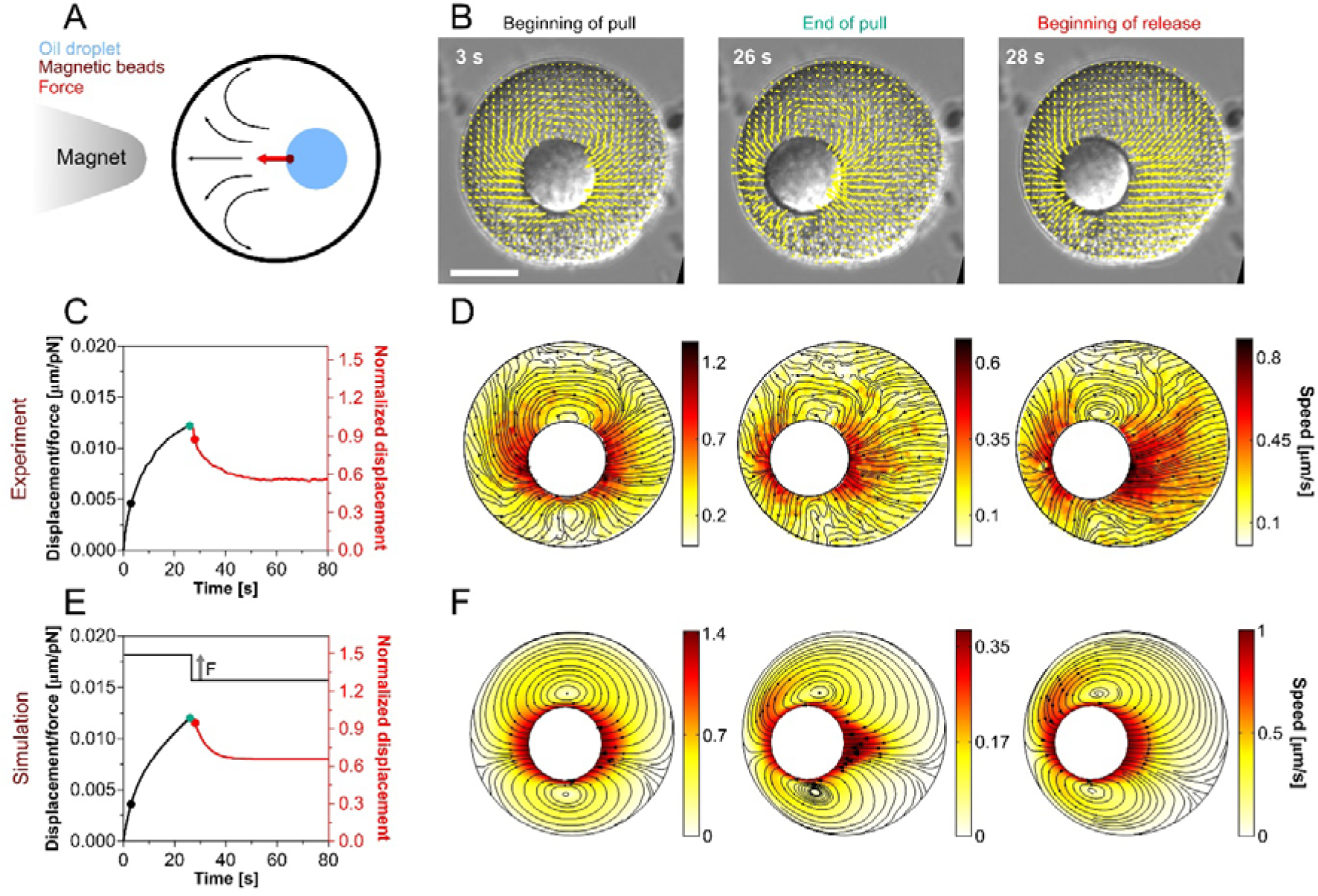
Experimental and theoretical mapping of shear cytoplasm flows created by the translation of a large object inside cells. (A) Schematic of a moving oil droplet and consequent viscoelastic flows. (B) Vector fields of cytoplasm flow obtained by PIV around a moving oil droplet at the beginning of the pulling phase, end of pulling, and beginning of the release phase overlaid on DIC images. (C) Scaled displacement curve of pulling phase and normalized curve of release phase for the same experiment as in B. Time points of temporal snapshots of the vector fields are indicated by color-coded circular markers on the curves. (D) Experimental streamlines and speed heat maps of cytoplasm flow for the same snapshots in B. (E) Scaled displacement curve of the pulling phase and normalized release phase obtained from finite-element simulations with input parameters taken from experimental measurements. (F) Numerical streamlines and speed heat maps of the same time points as in D. Arrowheads in the simulations are proportional to the speed. Scale bar 30 μm.

To test if experimental flows and viscoelastic responses could correspond to a simple interaction of the moving object with the surrounding cytoplasm fluid, we set up finite-element hydrodynamic simulations. We represented the oil droplet as a solid spherical object and displaced it with a constant force inside a confined Jeffreys’ viscoelastic medium (implemented as an equivalent Oldroyd-B model) and then released the force. As inputs of this model, we used the two viscosities and the second viscoelastic time-scale measured from experiments. The resulting simulations accounted for both flow fields organization and speed magnitude at various steps of the pulling and relaxation phases observed in experiments (Fig. 3F and Movie S3). We validated the simulations by computing the consistency between input and output values of material properties, which we measured from the simulated time-displacement curves following a 1D Jeffreys’ model, as in experiments. We found that the quantitative creep and recovery curves of the experiments were consistent with the simulation output, asserting that the 1D Jeffreys’ model was sufficient to describe the 3D viscoelastic response of the cytoplasm (Figs. 3E and S2B-F).

### Viscoelastic drag and restoring stiffness depend on confinement

Flow analysis suggested that the displacement of large objects generate shear flows from hydrodynamic interactions between the object and cell boundaries. We sought to test if these interactions could affect object mobility and notably enhance drag and/or restoring stiffness. For a sphere located at the exact center of a compartment filled with a Newtonian fluid, the viscous drag of the object is predicted to follow a modified Stokes’ law *γ_1_(λ)* = 6 *πηrC(λ)* where η is the fluid viscosity, r, the object radius and C a wall correction factor: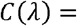 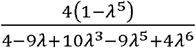 with λ =*r/R* the ratio of the sphere radius to that of the container (23). Therefore, such correction amounts to an increase in the effective viscosity felt by the object: *η*\**(λ)* = *ηC(λ)*. The correction factor asymptotically approaches 1 for small and weakly bounded object, is ~2.5 for objects ~30% of container size, reaches up to ~7 when objects are ~50% of container size, and diverges to infinity when object size approaches that of the container (Fig. 4A). By running 3D simulations, we confirmed the validity of this formula in the context of our particular pulling assay, and also predicted that confinement should have a similar impact on both viscous and elastic responses of the cytoplasm.

**Figure 4.**
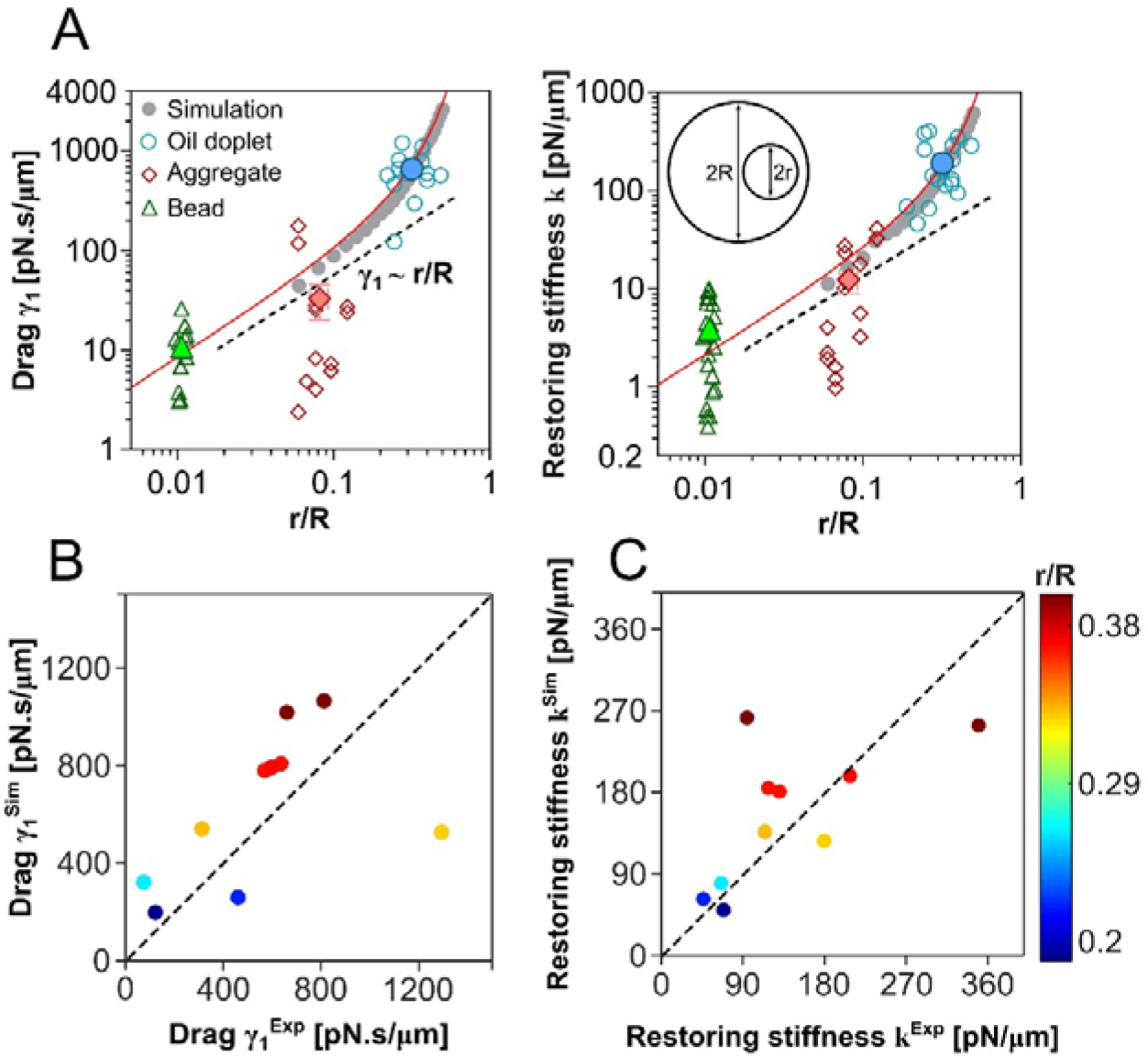
Cell confinement enhances cytoplasm viscoelastic resistance in a non-linear manner. (A) Viscous drag and restoring stiffness increase with the object size in a nonlinear manner, and deviate from linear Stokes’ law for confinement ratios larger than ~0.1. The red curve represents the analytical prediction for a newtonian fluid, gray solid circles are 3D simulations and solid colored symbols are average values for beads (n=21), aggregates (n=15), and oil droplets (n=18). (B-C) Individual viscous drags (B) and restoring stiffness (C) computed from experiments and simulations for individual droplets of various sizes in the cytoplasm. Dashed lines in B and C guide the eyes for a perfect match between experiment and models.

To test the relevance of confinement to experimental results, we next plotted measured viscous drags as a function of λ for the different sized objects (Fig. 4A). We found that experimental drags were in good agreement with the results of both analytical formula and 3D hydrodynamic simulations in the large intervals of confinement ratio. However, we noted some deviations from experiments for beads aggregates, in which drags were under theoretical curves. We interpret this deviation as a result of a plausibly more porous nature of aggregates and their non-perfect circular shapes. Importantly, for large oil droplets, the deviation from linear Stokes’ relationship became pronounced as the confinement ratio λ exceeded ~0.1-0.2, with correction factors that ranged from ~1 to ~9. In addition, as predicted by simulations, we found that the effect was similar for restoring stiffness, directly demonstrating that the proximity of boundaries can enhance both viscous and elastic cytoplasm resistance (Fig. 4A).

To better establish the relevance of this effect in dose-dependence, we next performed 3D simulations in the size range of spherical oil droplets, and compared predicted drags and restoring stiffness to individual experimental results. For this, we inputted average experimental values of cytoplasm viscosities and second viscoelastic time-scale, and varied droplet and egg size and initial position (see below) according to experiments. The agreement between experiments and simulations was in general very good, suggesting that experiments are close to the theoretical limit (Figs. 4B-C). Together, these results strongly suggest that hydrodynamic interactions between a large moving object and the cell surface, effectively enhances cytoplasm viscoelasticity at this scale.

### Slippage on boundaries reduces the effect of confinement on organelle mobility

As organelles and cell surfaces may come with different properties like rugosity or hydrophobicity, that may influence how the cytoplasm fluid interacts with these surfaces, we sought to evaluate the impact of stick or slip conditions on the confinement effects. In experiments, it was technically challenging to modulate slippage. Therefore, we performed a series of simulations to test how boundary conditions impact flow fields and size dependency. We considered three different scenarios: a no-slip boundary condition over the oil droplet and cortex surface, slippage only over the oil surface, and finally slippage over both oil and cortex surfaces. Interestingly, the general flow patterns were independent of boundary conditions chosen (Figs. 5A-C). However, the impact of dissipation for a sphere of a given size was reduced when the fluid slips on surfaces, yielding to a net increase in average fluid speed (Fig. 5D). Under the same applied force, the fluid speed drops as object size increases, but boundary conditions affect the speed in the same manner for spheres of different sizes (Fig. 5D). Accordingly, both viscous drag and restoring stiffness increased in a non-linear manner as a function of the confinement ratio λ under all boundary conditions, but the effect was more pronounced when the fluid could adhere to all surfaces (Figs. 5E-F). We conclude that the impact of confinement on organelle mobility may still be significant independently of the type of boundary conditions, but more pronounced for non-slip conditions.

**Figure 5.**
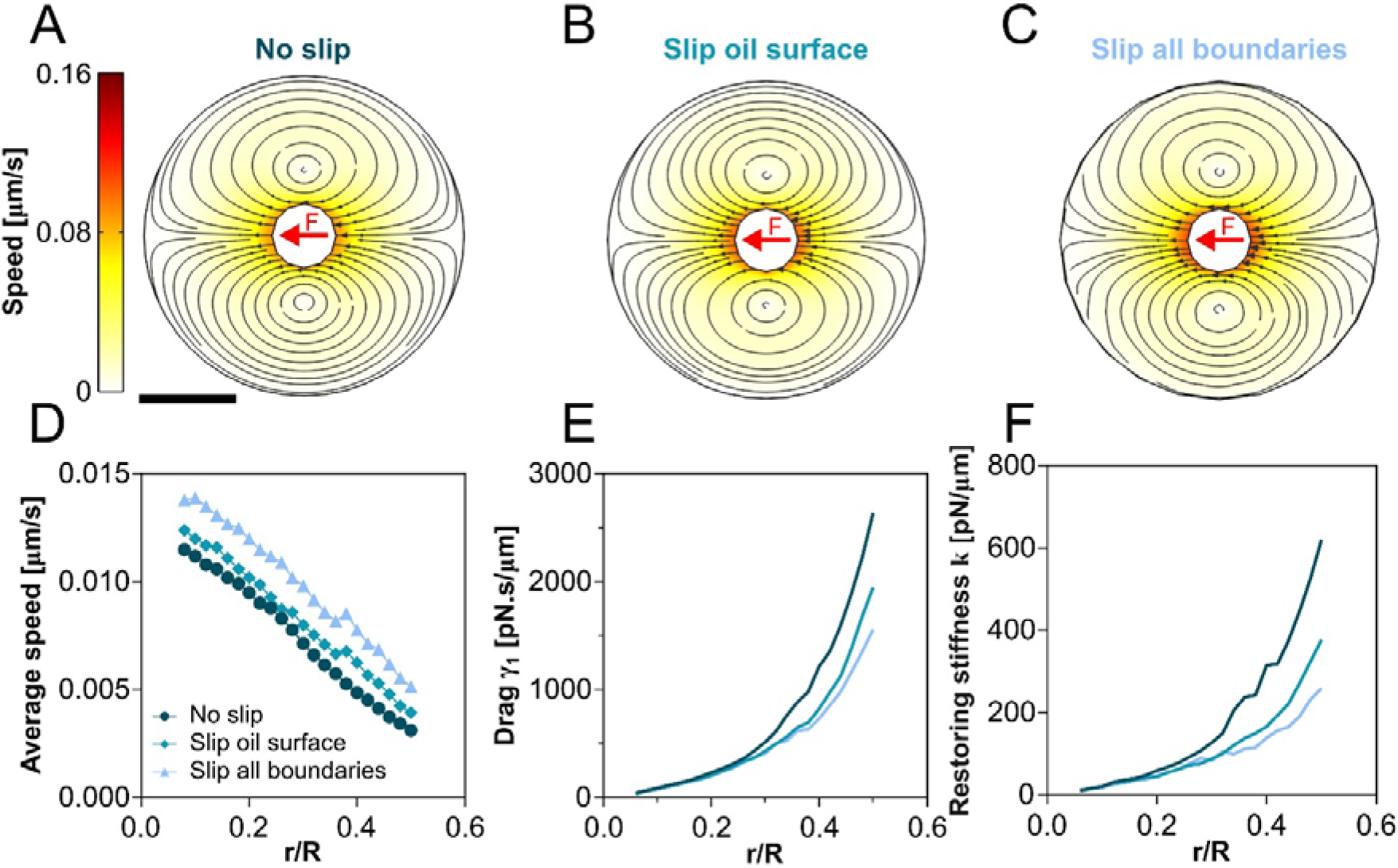
Impact of slip or stick boundary conditions for the influence of cell confinement on large object mobility. (A-C) Simulated streamlines and flow patterns of the cytoplasm for various boundary conditions on the oil droplet and cell surfaces. (D) Slippage over the boundaries increases the flow speed for all confinement ratio. The magnitude and direction of the pulling force are set to be the same for all spheres of different sizes. (E-F) Viscous drag and restoring stiffness grow in a non-linear manner for all types of boundary conditions, but the values of viscoelastic parameters are smaller when the fluid can slip over surfaces. Arrowheads in the simulations are proportional to the speed. Scale bar, 30 μm.

### Heterogeneity and anisotropy of drags and restoring stiffness

As organelles may adopt different locations within the cytoplasm, being *e.g*. initially closer to the cell surface, we sought to test if hydrodynamic interactions affect mechanical resistance of the cytoplasm depending on object’s initial position. We first performed a range of simulations. We considered a sphere with a confinement ratio of λ = 0.2 in a cell filled with Jeffreys’ fluid, and placed it at different positions along the force axis. We found that both drags and restoring stiffness increased as the object was closer to the cell surface, reaching an enhancement of ~2.5X at an offset distance of ~30% of cell size closer to the surface.

Simulating a much smaller sphere of λ = 0.04 as control, showed that this local effect is only relevant to relatively large objects (Figs. 6A-B). Interestingly, because of the linearity of the viscoelastic-model used, this position-dependent enhancement was perfectly symmetric, whether the object was pushed against or pulled away from the surface. Accordingly, analysis of fluid flows revealed symmetric flow maps with respect to the boundary, indicating a similar interaction of the fluid with the container surface in both cases (Fig. 6C). Finally, these symmetric enhanced effects were also observed for restoring stiffnesses (Fig. 6D). In a second sets of simulations, we also changed object position along an axis orthogonal to the force axis. In this situation, boundary conditions also enhanced cytoplasm resistance, but the effect was less pronounced than when objects moved orthogonal to the cell surface (Fig. 6A). Therefore, hydrodynamic interactions with cell boundaries may yield to anisotropic cytoplasm resistance facilitating the motion of large objects parallel as compared to orthogonal to the cell surface.

**Figure 6.**
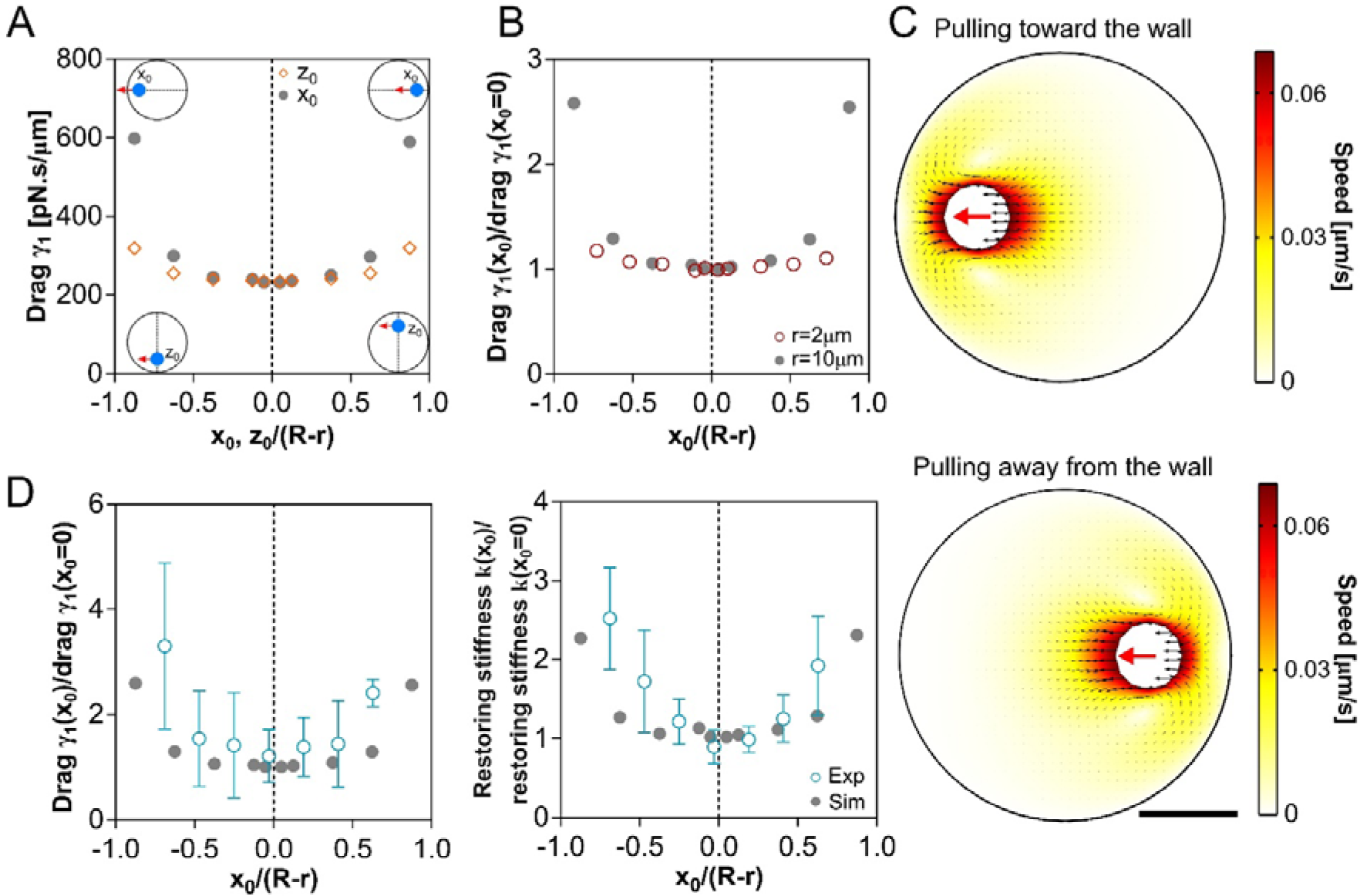
Cytoplasm viscoelastic properties are heterogeneous and anisotropic depending on object position. (A) Simulation of viscous drags experienced along the pulling force axis (x-axis) for objects positioned with offsets along the force axis or orthogonal to this axis (z-axis). (B) Simulations of viscous drags experienced by objects of bigger or smaller sizes. (C) Simulation of fluid hydrodynamics interaction with the cell surface as the sphere is pulled toward or away from the surface. (D) Measured viscous drags and restoring stiffness for oil droplets pulled along the x-axis starting from different initial positions (n=47 pulls, from 4 cells). The values of viscoelastic parameters have been renormalized to the values in the cell center, and experimental results are binned and overlaid with simulation results. Error bars correspond to the SD of data. Scale bar, 30 μm.

To test these predictions in real cells, we pulled oil droplets with different initial positions. To minimize variability arising from different droplets sizes, we performed successive pulls of the same droplets, spaced by sufficient time for the material to relax, and varied their initial position with magnetic tweezers. The average experimental values exhibited a minimum at the cell center, and increased in a symmetric manner as the droplet was placed closer to the cell surface, with a good alignment on simulations data (Fig. 6D). Interestingly, the position-dependent enhancement, for both drag and restoring stiffness was symmetric, as predicted from the linear model, suggesting that both cytoplasm compression and extension can result in viscous and elastic stresses. Together, these simulations and experiments demonstrate that cytoplasm viscoelasticity felt by large objects in a cell may be position-dependent and anisotropic.

## DISCUSSION

Here we used magnetic tweezers to displace large passive components of different sizes in the cytoplasm of living cells with unidirectional calibrated forces. Our results suggest that objects floating in bulk cytoplasm become coupled to the cell surface through the fluid, with no need for any direct cytoskeletal connections. This hydrodynamic coupling can occur over distances of up to tens of microns and enhance the viscoelastic resistance of the cytoplasm, thereby reducing organelle mobility. The strength of the coupling increases as the distance between the object and the surface reduces, so that larger components or components closer to the cell surface, become disproportionally harder to displace. Size- and position-dependency had previously been reported for the diffusion of smaller components in different cells. They were attributed to properties such as the hierarchy of cytoplasm pore sizes (33), glass-like transitions of bulk material (15), or to heterogeneities across cells (16). Our data for large objects, support a different physical origin, in which both size and position dependency can emerge solely from hydrodynamic interactions with the surface. As such, previously proposed alterations in bulk material properties or gradients in these properties, although not required here, could add up to the impact of hydrodynamic interactions in different cell types.

Our data suggest that the impact of hydrodynamic coupling becomes pronounced for objects over ~20-30% of cell size, which is the typical size ratio for organelles such as nuclei, mitotic spindles or microtubule asters. Accordingly, *in vivo* direct or indirect force measurements reported spindle, nuclei and aster drags that largely exceeded Stokes drags calculated from the size of these organelles and cytoplasm viscosity (20, 28, 34). Other relevant parameters that could impact cytoplasm resistance include organelle shape and porosity especially when considering fibrous-like structures such as asters and spindles (21, 35). Furthermore, as demonstrated here, the relative proximity of organelles to boundaries, may also greatly impact its mobility. As a consequence, we expect cell geometry to also influence organelle mobility. For instance, a nucleus moving in a tube-like cell such as fission yeast or a columnar epithelial cell (36, 37), is predicted to face very large resistance from the cytoplasm, given the little space left for the fluid to flow around the organelle (23). Although the role of the cytoplasm has been generally omitted in standard models for organelle positioning (7, 38), we anticipate they could impact our current appreciation of the mechanics of nuclear or spindle positioning, as well as that of shape changes.

Finally, our study also highlights large shear flows that form as a direct result of organelle motion under force, and cause the cytoplasm to recirculate over the scale of the whole cell (20). Cytoplasm flows can organize processes ranging from cortical polarity (39), RNA localization (40), to nuclear positioning and internal organization (41, 42). They typically emerge from cortical contractile acto-myosin flows that generate surface stresses that propagate through the cytoplasm (17, 18, 42, 43), or from the activity of bulk cytoskeleton and associated motors (44, 45). Based on our observations, we propose that the natural motion or rotation of large organelles during *e.g*. asymmetric division or nuclear translocation could create large scale flows, that may impact the polarization of both cytoplasmic and cortical elements. Further work on how cytoplasm and cortical material properties and forces are integrated to control organelle mobility, will enlarge our understanding of cellular organization.

## ACKNOWLEDGMENT

We thank all members of the Minc lab for discussion and technical help. J.N. is supported by a fellowship from the ARC foundation (PDF20191209818) and is an EMBO Non-Stipendiary Fellow (ALTF 881-2019). S.D. acknowledges support from ITMO Cancer of Aviesan on funds managed by Inserm. This work was supported by the Centre National de la Recherche Scientifique (CNRS), the Université Paris Cité, and grants from La Ligue Contre le Cancer (EL2021.LNCC/ NiM), the Agence Nationale pour la Recherche (ANR, “TiMecaDev”), the Fondation Bettencourt Schueller (“Coup d’élan”), and the European Research Council (ERC CoG “Forcaster” no. 647073) to N.M.

## Material and methods

### Sea urchin gametes

Purple sea urchins (*Paracentrotus lividus*) were obtained from Roscoff Marine station (France) and maintained at 16 °C in aquariums of artificial seawater (ASW; Reef Crystals, Instant Ocean). Gametes were collected by intracoelomic injection of 0.5 M KCl. Eggs were rinsed twice with ASW, kept at 16 °C, and used on the day of collection.

### Injection

Unfertilized eggs were placed on protamine-coated 50 mm glass-bottom dishes (MatTek Corporation) after removing the jelly coat through an 80-μm Nitex mesh (Genesee Scientific). The bead suspensions were injected using a micro-injection system (FemtoJet 4; Eppendorf) and a micro-manipulator (Injectman 4; Eppendorf). Injection pipettes were prepared from siliconized (Sigmacote) borosilicate glass capillaries (1 mm diameter). Glass capillaries were pulled using a needle puller (P-1000; Sutter Instrument) and ground with a 40° angle on a diamond grinder (EG-40; Narishige) to obtain a 10 μm aperture. Injection pipettes were back-loaded with ~2 μl of bead suspension before each experiment, and were not re-used.

### Immunostaining

Immunostaining was performed using procedures described previously (46). Samples were fixed for 70 minutes in 100 mM Hepes, pH 6.9, 50 mM EGTA, 10 mM MgSO4, 2% formaldehyde, 0.2% glutaraldehyde, 0.2% Triton X-100, and 400 mM glucose. To reduce autofluorescence, eggs were then rinsed in PBS and placed in 0.1% NaBH4 in PBS freshly prepared, for 30 min. Eggs were rinsed with PBS and PBT (PBS + 0.1% TritonX) and blocked in PBT supplemented with 5% goat serum and 0.1% bovine serum albumin (BSA) for 30 minutes. Samples were rinsed with PBT before adding primary antibodies. For microtubule staining, cells were incubated for 48 h at 4 °C with a mouse anti-α-tubulin (DM1A; Sigma-Aldrich) primary antibody at 1:5,000 in PBT, rinsed 3 times in PBT and incubated for 4 hours at room temperature with anti-mouse secondary antibody coupled to Dylight 488 (Thermo Fisher Scientific) at 1:1,000 in PBT for 4-5 hours. To stain F-actin, samples were also incubated together with secondary antibodies in a solution of Rhodamine Phalloidin at 4 U/ml in PBT. Eggs were washed three times in PBT then twice in PBS, transferred in 50% glycerol in PBS, and finally transferred into mounting medium (90% glycerol and 0.5% N-propyl gallate in PBS).

### Hypoosmotic and hyperosmotic shocks

To elevate cytoplasm crowding with hyperosmotic shocks, ASW was prepared to 120% of its normal content by adjusting the amount of DI water to salts mixtures. The eggs laid down on protamine-coated dishes inside the hyperosmotic water and then injected. Higher concentrations of 150% were tested, but droplets could not be pulled in such cytoplasm. To reduce cytoplasm density with hypoosmotic shocks, the eggs were first injected and then ASW was diluted by the same volume of DI water equal to ASW inside the dishes, to reach an ASW at 50% of its normal content.

### Magnet force calibration

Magnetic forces were calibrated *in vitro* following procedures described previously (20, 28, 47). The magnetic force field created by the magnet tip was first characterized by pulling super-paramagnetic 1 μm Dynabeads (MyOne Streptavidin C1; Thermofisher) in a viscous test fluid (80% glycerol; viscosity 8.0×10^−2^ Pa.s at 22 °C) along the principal axis of the magnet tip. Small motion of the fluid was subtracted by tracking 0.5 μm non-magnetic fluorescent tracers (Molecular probes; Invitrogen) in the same suspension. The speed of magnetic beads V was computed as a function of the distance to the magnet, to obtain and trace the decay function of the magnetic force, which was fitted using a double exponential function (Fig. S1E).

To calculate the dependence of the force on aggregate size, bead aggregates from the same beads as those used *in vivo* in sizes ranging from 2 to 7 μm, similar to that observed in oil droplets, were pulled in the same fluid as above. The speed V_a_ was measured and transformed into a force using Stokes’ law F = 6πηRV_a_, where η is the viscosity of the test fluid, R the aggregate effective radius defined as the geometric mean of the longest and shortest axes of the aggregate 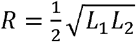. In live cell experiments the size of the aggregates inside the oil droplets was measured at three different positions in the bright field and averaged (Fig. S1D). The force–size relationship at a fixed distance from the magnet was well represented and fitted by a cubic function (Fig. S1F). These speed-distance and force-size relationships were combined to compute the magnetic forces applied to the beads and aggregates as a function of time, from the size of aggregates and their distance to the magnet tip.

### Magnetic force application

Magnetic tweezers were implemented as described previously (20, 28, 47). The magnet probe used for force applications *in vivo* was built from three rod-shaped strong neodymium magnets (diameter 4 mm; height 10 mm; S-04-10-AN; Supermagnet) prolonged by a sharpened steel piece with a tip radius of ~60 μm to create a magnetic gradient. The surface of the steel tip was electro-coated with gold to prevent oxidation. The probe was controlled with a micromanipulator (Injectman 4; Eppendorf) and mounted on an inverted epifluorescent microscope.

Super-paramagnetic 1 μm Dynabeads (MyOne Streptavidin C1; Thermofisher) were pulled inside the cytoplasm as single beads. To prepare beads for injection, a 1 μl bead suspension was diluted in 100 μl washing solution (1 M NaCl with 1% Tween-20) and sonicated for 20 minutes. Beads were then separated by a magnet, rinsed in 100 μl PBS BSA 1%, and sonicated for 20 minutes after 5 minutes of incubation. The beads were then re-suspended in 20 μl of 2 μg/ml Atto565-biotin (Sigma-Aldrich), incubated for 5 minutes, and finally re-suspended in 20 μl PBS.

Super-paramagnetic 800 nm particles (NanoLink; Solulink) were used from their propensity to form large aggregates inside the eggs. A solution of 10 μl of undiluted streptavidin beads was first washed in 100 μl of washing solution and sonicated for 1 hour. The beads were then incubated for 15 minutes in 100 μl of 2 μg/ml Atto565-biotin (Sigma-Aldrich), rinsed in 100 μl PBS, and finally re-suspended in 20 μl. To form aggregates in cells, beads first were pulled by the magnet on one side of the egg to form an aggregate close to the cortex gathering all the beads inside the cytoplasm. Then, the dish was rotated and the aggregate was pulled from the other side of the egg. The aggregates often stretched along the pulling direction which could potentially reduce their viscous drag.

To pull oil droplets in the cytoplasm, a suspension of 10 μl of 1.2 μm hydrophobic superparamagnetic beads (magtivio; MagSi-proteomics C18) was washed in 100 μl of 30, 50, and 70 percent ethanol solution. It was then dried in vacuum for ~20 minutes and re-suspended in ~10 μl soybean oil (Naissance; Huile de soja). All the bead suspensions were kept at 4 °C until use. All oil-injected eggs in each sample were surveyed to identify oil droplets with a sufficient amount of beads needed for pulling. After approaching the magnet, beads inside the oil progressively formed an aggregate and slowly moved toward the magnet while the oil droplet was stationary until the aggregate contacted the oil-cytoplasm interface on the side facing the magnet tip.

Large aggregates could not usually cross the interface due to the oil surface tension, aggregate size, and hydrophobic properties of the beads, and were used to pull oil droplets in the cytoplasm. Upon force application, the magnet was quickly retracted when the oil distance to the cell cortex was ~10 μm, which usually caused aggregate detachment from the oil-cytoplasm interface and backward relaxation of the droplet along the initial pulling force axis. The initial position-dependent experiments with the oil droplets were done similarly as for aggregates. Oil droplets were first pulled toward the cortex on one side of the egg. However, the droplets could not stay very close to the cortex because of the viscoelastic response of the cytoplasm. Then, the dish was rotated and the oil droplet was pulled from the opposite side along the diagonal direction. Droplets were displaced slowly to change the offset and suppress the oil droplet from recoiling toward its initial position (20). The droplet was pulled again in the same direction after at least a 5 minutes delay to ensure a complete relaxation of the droplet.

### Imaging

Time-lapses of oil droplets, aggregates, and beads moving under magnetic force were recorded on two inverted microscope set-ups equipped with a micromanipulator for magnetic tweezers, at a stabilized room temperature (18-20 °C). The first set-up was an inverted epifluorescence microscope (TI-Eclipse; Nikon) combined with a complementary metal-oxide-semiconductor (CMOS) camera (Hamamatsu), using a 20X DIC dry objective (Apo; NA 0.75; Nikon) and a 1.5X magnifier, yielding a pixel size of 0.216 μm. The second one was a Leica DMI6000 B microscope equipped with an A-Plan 40x/0.65 PH2 objective yielding a pixel size of 0.229 μm, equipped with an ORCA-Flash4.0LT Hamamatsu camera. Both microscopes were operated with Micro-Manager (Open Imaging). Imaging was done in DIC/fluorescence/bright field at a rate of 1 frame per s. Immuno-stained eggs presented in Figs. S1A-B were imaged on a spinning-disk confocal microscope (TI-Eclipse; Nikon) equipped with a Yokogawa CSU-X1FW spinning head, and a Prime BSI camera (Photometrics), using a 60X water-immersion objective (Apo; NA 1.2; Nikon).

### Tracking bead and droplet positions

Magnet tip position was recorded in the bright field or fluorescent images through the fluorescent beads attracted onto the magnet. Oil droplets and aggregates time-lapse images were rotated to align their displacement vector antiparallel to the horizontal x-axis meaning that they all move from the right to the left. The position of beads and aggregates were tracked from their fluorescence signal using the TrackMate plugin in Fiji. The trajectories of the beads were rotated to align their displacement from the right to the left. Bright-field images of oil droplets were segmented in Fiji from the contrast at the periphery of the droplet and tracked using the TrackMate plugin. Displacement of the oil droplet was corrected when the egg had moved during the pulling.

### Viscoelastic parameter calculation

Oil droplet and bead displacements were fitted with a Jeffreys’ model using a custom written code in Matlab (Mathwork) to compute viscoelastic parameters (Figs. 1B-C). For the rising phase, the position was fitted using:

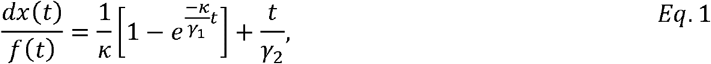

where *dx* is the displacement along the x-axis and *f* is the magnetic force. This re-scaling of the displacement by force allows to compensate for variations in force amplitude during each pull, and implicates that we assume that viscoelastic responses are mostly linear. These fits allowed to compute the restoring stiffness, *κ*, and the viscoelastic drags *γ*_1_ and *γ*_2_ on oil droplets and beads, allowing to compute viscoelastic time-scales as *τ*_1,2_ = *γ*_1,2_/κ.

Trajectories of beads during the relaxation phase were influenced by the intrinsic cytoplasm fluctuations and beads started random motions after moving a few steps backward (Fig. 1A). It was assumed the viscoelastic response of beads was ended and that the Brownian motion started when the angle between two successive steps exceeded 90 degrees. Relaxation phases of beads, aggregates, and oil droplets were fitted using:

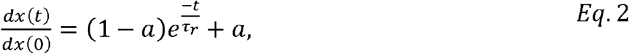

where t = 0 corresponds to the time of the end of force application, to compute the decay time-scale τ_r_.

Fit to the data were obtained for each single experiment using optimization method in Matlab in which the results are less sensitive than Nonlinear least squares method to the initial values.

### Flow analysis

The recorded oil droplet images in DIC/bright field were analyzed using the particle image velocimetry PIVlab tool in Matlab. The exterior of the egg and oil droplet surface at each frame were masked to be excluded from the analysis. Contrast limited adaptive histogram equalization (CLAHE) and two-dimensional Wiener filter with accordingly windows of 20 and 3 pixels widths were applied on the images in the pre-processing steps for noise reduction. Image sequences were investigated in the Fourier space by four interrogation windows with 64, 32, 16 and 8 pixels widths and 50% overlapped area. The spline method was used for the window deformation and subpixel resolution obtained by two-dimensional Gaussian fits. The distribution of velocity components of vectors for each set was visually inspected and restricted to remove outliers in the post-processing stage. Moreover, two other filters based on the standard deviation and local median of velocity vectors were applied to validate the vector fields. The output vector fields after smoothing were used for further analysis and plotting flow maps and streamlines in MATLAB.

### Finite element hydrodynamic simulations using COMSOL

Pulling of the object in the cytoplasm was modeled as a time dependent problem using the finite element software COMSOL, using an Oldroyd-B viscoelastic model. Oldroyd-B is a three element viscoelastic fluid model consisting of a Maxwell element in parallel with a viscous element represented by the constitutive equation, 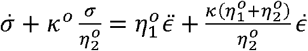 where σ and □ denote stress and strain and dot represents temporal derivative (48). The parameters κ°, η°_2_, and η°_1_ respectively indicate stiffness, viscosity of the dashpot in series with the spring, and viscosity of the dashpot parallel to the spring. This viscoelastic model is equivalent to Jeffreys’ model as one can write the Oldroyd-B model parameters as a function of the Jeffreys’ parameters measured experimentally:

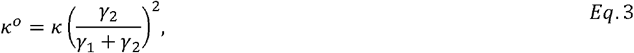

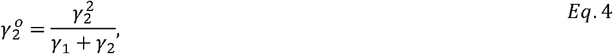

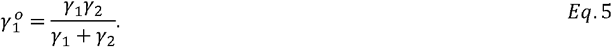

The viscous drag γ_2_° is associated to the dashpot in series with the elastic element describing the reorganization of cytoplasm. The drag γ_1_° is connected to the dashpot parallel to the elastic element serving for the viscosity of cytoplasm, and κ° is the cytoplasm stiffness. We could obtain the viscosity through the Stokes’ law γ = 6πηr. However, we also had to take into account the correction factor from confinement *C(λ)*, yielding γ(λ) = 6πηrC(λ), with:

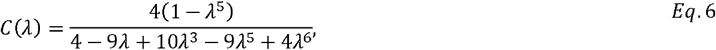

where λ = r/R was the ratio of oil droplet radius to egg radius. Considering the simulation results (Fig. S3B), we assumed the viscoelastic time-scale τ_1_ = γ_1_/κ as a constant material property which allowed us to apply the same confinement correction factor *C(*λ) for calculating stiffness inside the cell.

For simplicity, we modeled the oil droplet as a spherical elastic sphere with different radii r, and set its initial position in the center of the coordinate system except for the one-by-one and position dependent simulations (Figs. 3E-F, 4-6, and S2). The egg was represented as a large non-deformable sphere of radius R. The cytoplasm was interacting with the pulled object by including viscoelastic flow and solid mechanics modules in COMSOL through stresses at the boundaries. A Heaviside step function was used to apply a constant force as a body load with an amplitude equals to the average magnetic force in the experiments and the same pulling time. A no-slip wall condition was set as a default on the boundaries and changed to slip condition on the walls of droplet and cell to examine its impact on the results (Fig. 5).

We first verified the lack of impact of the elasticity of the pulled objects by varying this parameter over a very wide range. The elasticity had an overall little impact, lesser than 20%, on the viscosities. The viscoelastic time-scale τ_2_ was differing from the actual cytoplasm properties by less than ~44% (Fig. S2B). As COMSOL is a finite-element simulation software in which elements are discretized by a mesh, we also tested several mesh sizes including fine, normal, coarse, and coarser and observed the results from different mesh size were not significantly different (Fig. S2C). The simulations were performed using a coarse mesh size and the size of the pulled object was reduced to the extent that re-meshing was feasible for the combination of used parameters and simulation time was reasonable. We then needed to ensure that bulk viscosities and viscoelastic time-scales, τ_2_ = γ_2_/κ could be correctly evaluated from the drags and spring constants by fitting the one-dimensional Jeffreys’ model to the simulated displacement curves. In the COMSOL model, we varied input cytoplasm properties η_1_, η_2_, and τ_2_ around the average experimental values and measured the simulated displacements. We next fitted the one-dimensional Jeffreys’ model to the scaled pulling curves, extracted and converted the output bulk properties to Oldroyd-B model which were close to the input values (Figs. S2D-F). The maximum deviation between the input and output viscosities in the range of values that were tried was less than ~20%. These results show that the one-dimensional model is sufficient to describe bulk cytoplasm properties in three dimensions.

Heat maps, flow fields, and streamlines of the simulations were plotted using COMSOL and the rest of the analysis were done using custom written codes in MATLAB similar to the experimental data. The input parameters of the simulations for the Figs. 4-6 are summarized in Table S1.

## Supplemental Information

**Table S1.**
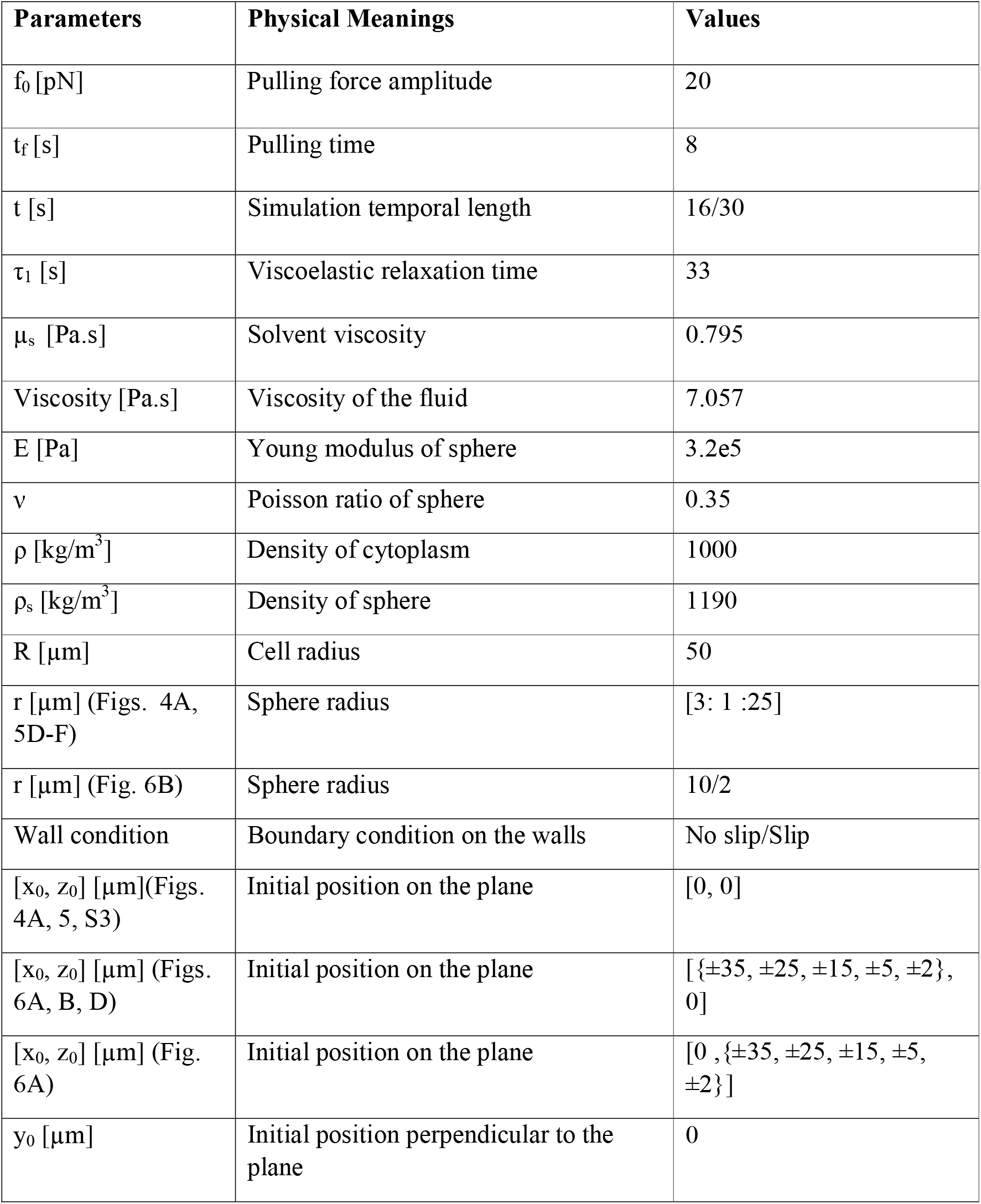
Parameters used in 3D finite-element simulations.

**Figure S1.**
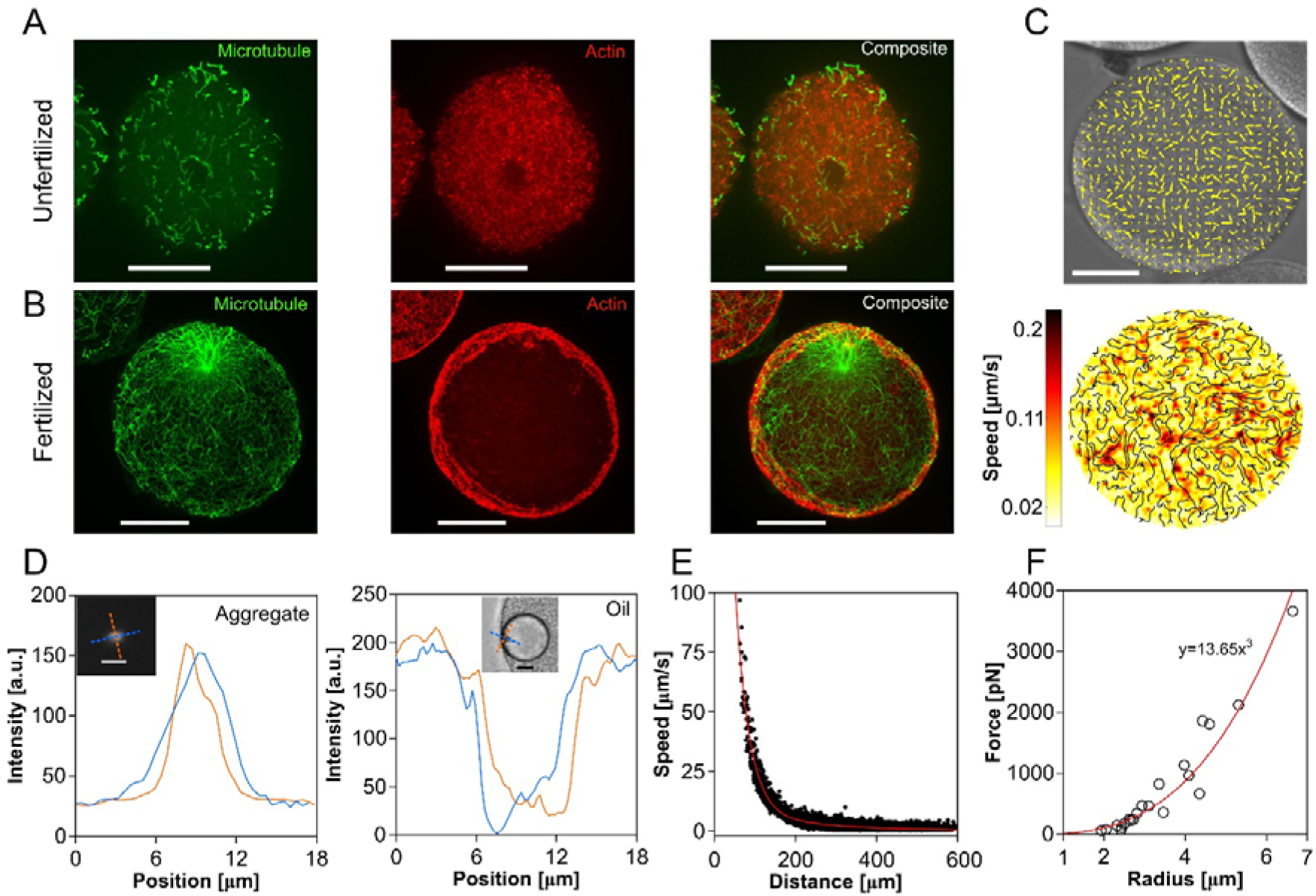
Cytoplasm organization in unfertilized eggs, and magnetic force calibration. (A) Confocal images of an unfertilized sea urchin egg fixed and stained for Microtubules and F-actin. (B) Confocal images of a fertilized sea urchin egg fixed and stained for MTs and F-actin. Images are maximum intensity projections of a mid-slice egg stack. (C, top) Vector field for internal flows in an unfertilized egg overlaid on DIC images. (Bottom) Heat map of the flow speed and streamlines for the same frame as above. Scale bars in A-C, 30 μm. (D) Intensity profile along the two main axes of hydrophilic bead aggregate in eggs imaged in fluorescence, and of hydrophobic bead aggregate inside the oil droplet imaged in bright field. These profiles are used to compute the mean size of aggregates. Scale bars in the inset, 10 μm. (E) Speed of 1 μm magnetic beads moved in a test viscous fluid, plotted as a function of their distance to the magnet tip, used to calibrate the magnet. The red curve is a double exponential fit. (F) Magnetic force at 70 μm from the magnet tip, plotted as a function of aggregate size to calibrate how magnetic forces evolve as a function of aggregate size. The red curve is a cubic fit.

**Figure S2.**
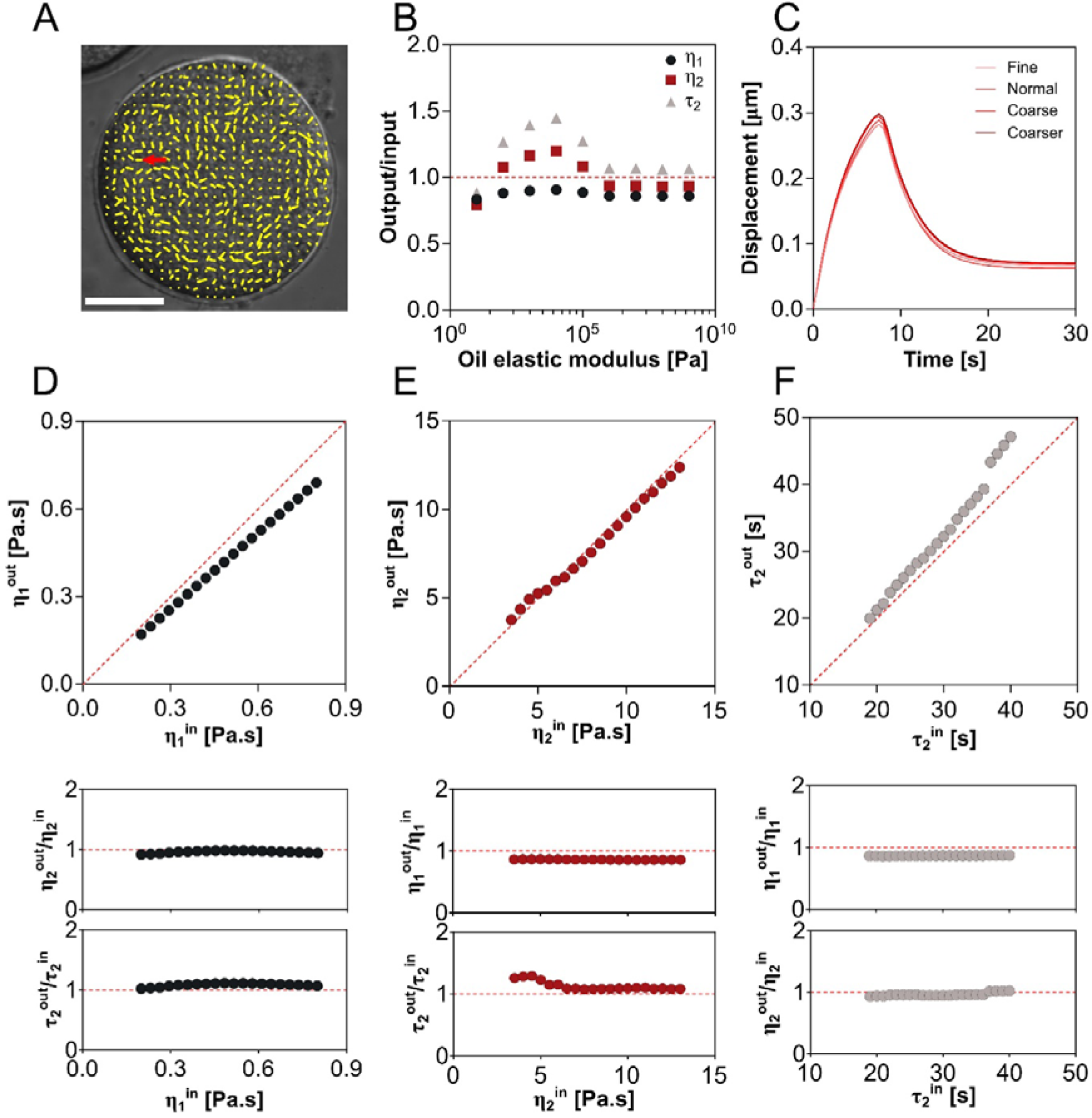
Validation of finite element 3D hydrodynamic simulations. (A) Vector field of cytoplasm flows as a 1 μm bead is pulled through the cell. The moving direction of the bead is indicated by the red arrow. (B) Normalized simulation output values of the two viscosities and time-scale in the Oldroyd-B model plotted as a functions of oil elastic modulus. (C) Displacement curve of pulling and release phase for the same sphere of 20 μm in diameter using different discretization mesh sizes in the simulation. (D, top) Output values of viscosity η_1_ computed from the 3D simulation results using the Jeffreys’ model plotted as a function of the input value given to the simulation. (Middle) Plot of the output/input ratio of the second viscosity η_2_ in the Oldroyd-B model as a function of different input values for η_1._ (Bottom) Plot of the output/input ratio of the time-scale τ_2_ =k/η_2_ in the Oldroyd-B model as a function of different input values for η_1._ (E) Output values of η_2_, output/input ratio for η_1_, and output/input ratio of τ_2_ plotted as functions of input values for η_2_. (F) Output values of τ_2_, output/input ratio for η_1_, and output/input ratio of η_2_ plotted as functions of input values for τ_2_. Dashed red lines guide the eyes and correspond to equal values for input and output. Scale bar, 30 μm.

**Figure S3.**
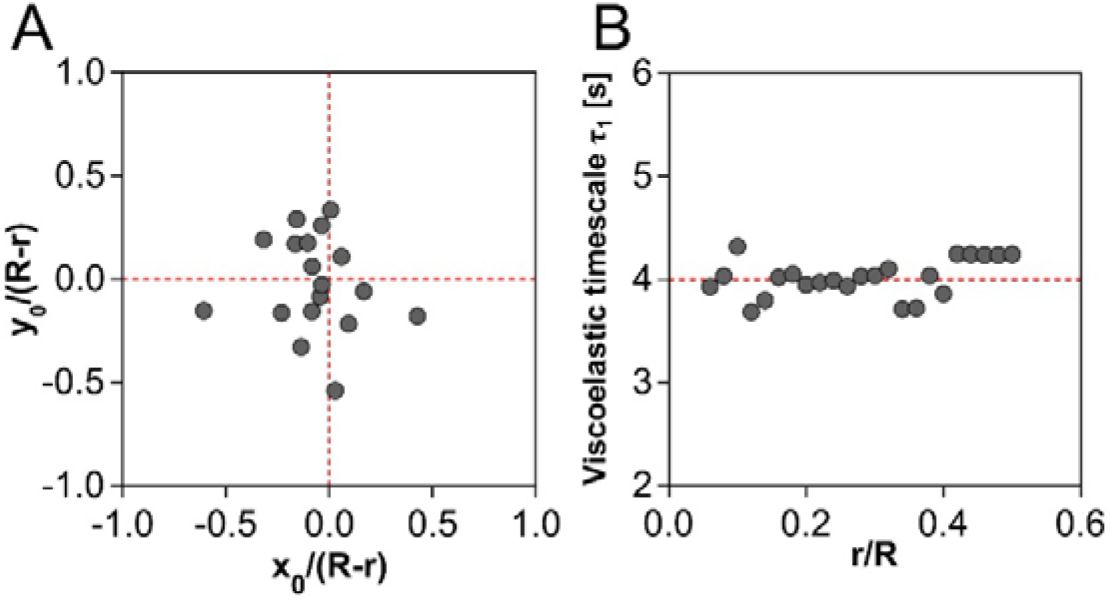
Initial position of droplets and viscoelastic time-scales at different aspect ratio. (A) Distribution of normalized initial positions of oil droplets in experiments presented in Fig. 4. (B) Viscoelastic time-scale computed from the simulation as a function of confinement ratio.

## Supplemental Movie Legends

**Movie S1. Pulling objects at different length scales in the cytoplasm**. Time-lapses of pulled objects of diverse sizes using calibrated magnetic forces, directed from right to left in unfertilized sea urchin eggs. Time is in second and the scale bar is 30 microns.

**Movie S2. Magnetized oil droplets pulled and recoiled at various crowding conditions of cytoplasm**. Time-lapses of eggs at distinct crowding conditions injected with oil droplets containing magnetic beads, pulled with magnetic tweezers, and let to recoil. Time is in second and the scale bar is 30 microns.

**Movie S3. Experimental and numerical mapping of cytoplasm flows created by the translation of a large object in cells**. The Vector field of cytoplasm flows overlaid on experimental images as a large object was moved in the cell. Streamlines and speeds were compared between the experiment and COMSOL simulation with the same parameters as that of the experiment. Time is in second and the scale bar is 30 microns.

